# AAV-only targeting of ventral tegmental area dopamine neurons for optical self-stimulation studies in mice

**DOI:** 10.64898/2026.02.10.705134

**Authors:** Ezequiel Marron Fernandez de Velasco, John C. Brent, Alex L. Welter, Praseuth Yang, Annelise Wickman, Eric H. Mitten, Kevin Wickman

**Author notes:** <u>co-corresponding authors</u> 6-120 Jackson Hall, 321 Church Street SE, Minneapolis, MN 55455.

## Abstract

Studies employing optogenetic approaches in rodent models have highlighted the important contribution of ventral tegmental area (VTA) dopamine (DA) neurons to reward, learning, and motivation. Selective manipulation of VTA DA neurons is generally achieved in these studies using transgenic mouse or rat lines that express Cre recombinase under the control of a promoter active in DA neurons, combined with intra-VTA infusion of adeno-associated virus (AAV) vectors harboring Cre recombinase-dependent expression cassettes. Reliance on transgenic Cre driver lines is expensive and decreases study efficiency, and available driver lines have unique limitations. Here, we report the development of an AAV-only approach that permits genetic access to VTA DA neurons and can support optogenetic self-stimulation in mice. We used a 2.5 kb fragment of the mouse tyrosine hydroxylase promoter (mTH) to drive Cre expression in VTA DA neurons. Intra-VTA co-infusion of AAV8-mTH-Cre with an AAV vector harboring a Cre-dependent yellow fluorescent protein expression cassette yielded high efficiency (82%) and high fidelity (73%) targeting of tyrosine hydroxylase-positive VTA neurons in C57BL/6J mice. Co-infusion of AAV8-mTH-Cre with a vector harboring a Cre-dependent channelrhodopsin (ChR2) expression cassette permitted optical regulation of VTA neurons with electrophysiological features consistent with VTA DA neurons. Moreover, C57BL/6J mice expressing ChR2 in VTA DA neurons rapidly acquired optical self-stimulation behavior. Thus, this AAV-only approach should facilitate investigation of VTA DA neuron contributions to reward-related behaviors and permit comparative assessments in reward circuit function in inbred and mutant mouse strains.

## INTRODUCTION

Drugs of abuse share an ability to enhance dopamine (DA) transmission in the mesocorticolimbic system [1], an interconnected network of brain regions including the nucleus accumbens (NAc) and medial prefrontal cortex (mPFC) that mediates the processing and learning related to rewarding stimuli [2]. DA neurons of the ventral tegmental area (VTA) are an integral component of the mesocorticolimbic system [3-5]. Optogenetic manipulation of VTA DA neurons is a powerful means to assess the contribution of these neurons to reward-related behavior [5,6]. For example, optogenetic stimulation of VTA DA neurons enhances locomotor activity in mice [7-10], mimicking the unconditioned DA-dependent response evoked by drugs of abuse. Optogenetic stimulation of VTA DA neurons can induce a conditioned place preference [7].

Studies employing optogenetic tools have shown that rodents will engage in instrumental responding to self-stimulate VTA DA neurons [8,11]. Optical self-stimulation (OSS) of VTA DA neurons allows for response-contingent activation of VTA DA neurons, mimicking the ability of addictive drugs to enhance DA transmission in the mesocorticolimbic system. OSS of VTA DA neurons is sufficient to induce key cellular and behavioral hallmarks of repeated drug exposure in mice, including potentiation of glutamatergic neurotransmission in the NAc and resistance of reward-seeking behavior to punishment [12]. OSS has also highlighted critical roles of discrete VTA DA neuron projections in reinforcement and cue-induced reinstatement of reward-seeking behavior [13,14].

Genetic manipulation of VTA DA neurons in support of OSS and other addiction-relevant paradigms generally involves a transgenic rodent line expressing Cre recombinase under the control of the promoter for tyrosine hydroxylase (TH) [15,16] or the DA transporter (DAT) [17,18], together with intra-VTA infusion of Cre-dependent adeno-associated virus (AAV) vector [19,20]. The use of transgenic Cre driver lines for studies of VTA DA neurons limits investigation to genetically tractable species and comes with the associated expense of procuring and maintaining mutant strains. These costs are compounded if breeding the Cre driver line with a particular inbred or mutant strain (*e*.*g*., a knockout mouse line) is desired.

Transgenic Cre driver lines also have unique limitations. In TH-Cre knock-in mice, for example, Cre-dependent transgene expression has been reported in a significant fraction of neurons within and surrounding the VTA that exhibit little-or-no TH labeling [19,20], including a unique population of VTA neurons that project to the lateral habenula and release GABA but not DA [21]. DAT-Cre mice are thought to permit more selective targeting of VTA DA neurons [19], but some TH-positive VTA neurons exhibit little-or-no DAT expression [22-25]. Thus, studies employing DAT-Cre mice may miss the critical contributions of some VTA DA sub-populations to reward-related behaviors. Genomic insertion of the Cre recombinase transgene can also influence endogenous gene expression. Indeed, a commonly used DAT-Cre knock-in mouse line (*Slc6a3*^*tm1*.*1(cre)Bkmn*^/J) exhibits reduced DAT protein levels in the striatum, ranging from 17% reduction in hemizygous animals to a 47% reduction in homozygous mutants [18], correlating with reduced DA reuptake capacity and aberrant reward-related behaviors [26-28].

Given the various limitations associated with the use of transgenic Cre driver lines, we sought to develop an AAV-only approach to VTA DA neuron manipulation that could support OSS studies in any mouse strain. Our approach exploits a relatively short fragment (2571 bp) of the mouse TH (mTH) promoter [29]. We demonstrate that the mTH promoter supports Cre-dependent transgene expression in VTA DA neurons in C57BL/6J mice with a transduction efficiency and targeting fidelity comparable to that reported for TH-Cre knock-in mice. We further show that this AAV-only approach can be used for optogenetic regulation of VTA DA neurons and OSS studies in C57BL/6J mice.

## RESULTS

### AAV-only targeting of VTA dopamine neurons in C57BL/6J mice

To gain genetic access to VTA DA neurons in C57BL/6J mice, we used a relatively short (2571 kb) fragment of the mouse TH promoter (mTH). Intravenous administration of an AAV-PHP.eB vector harboring this promoter yielded high efficiency (86%) and high fidelity (81%) reporter expression in TH-positive neurons in the mouse VTA [29]. Given the genetic payload limitations of AAV [30], we used a dual-virus approach to promote selective transgene expression in VTA DA neurons. One vector included the coding sequence for Cre recombinase positioned downstream of the mTH promoter (AAV8-mTH-Cre). The other vector harbored a Cre-dependent cassette for EYFP (AAV8-EF1α-DIO-EYFP). A similar dual-virus strategy involving the rat TH promoter has been used to achieve selective VTA DA neuron manipulation in rats and mice [31,32].

C57BL/6J mice received an intra-VTA infusion of a 1:1 mixture of AAV8-mTH-Cre and AAV8-EF1α-DIO-EYFP (**Fig. 1A**). We targeted the parabrachial pigmented nucleus of the VTA (PBP) to minimize off-target labeling of DA neurons of the substantia nigra. After a 2-3-wk incubation period, we used immunohistochemistry to quantify the efficiency and fidelity of viral targeting of TH-positive neurons in the VTA. We detected EYFP expression throughout the VTA of C57BL/6J mice that received the virus mixture (**Fig. 1B**), but not in mice that received either vector alone (not shown). Within the region of viral transduction, 82.0±0.4% of TH-positive neurons expressed EYFP and 73.1±3.1% of EYFP-positive neurons expressed TH (**Fig. 1B**,**C**).

**Figure 1.**
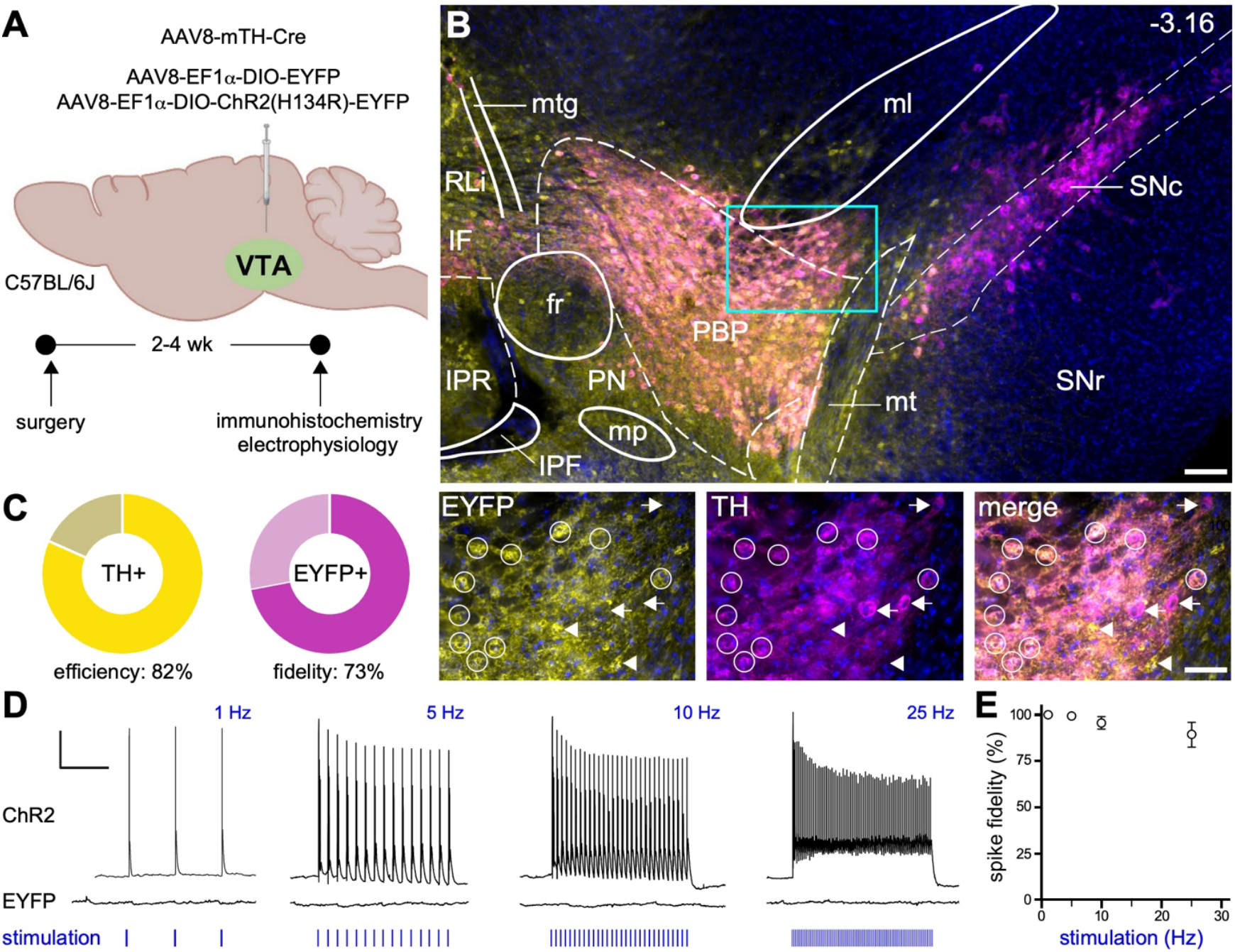
AAV-only genetic manipulation of VTA DA neurons in C57BL/6J mice. **A**. Male and female C57BL/6J mice were given intra-VTA infusions of AAV8-mTH-Cre and either AAV8-EF1α-DIO-hChR2(H134R)-EYFP or AAV8-EF1α-DIO-EYFP. Following a 2-4 wk recovery and incubation period, mice were used in immunohistochemistry or slice electrophysiology experiments. **B**. Fluorescent image showing a coronal section containing the VTA (-3.16 mm from bregma) of a mouse treated with intra-VTA AAV8-mTH-Cre and AAV8-EF1α-DIO-EYFP. EYFP fluorescence is denoted in yellow, TH immunolabeling is shown in fuchsia, and nuclear (DAPI) labeling is shown in blue; scale: 100 μm. The 3 images below show separate EYFP and TH labeling, and the image overlay (merge), within the region defined by the blue rectangle; scale: 50 μm. White circles highlight cell bodies with EYFP and TH labeling. White arrows highlight cell bodies with TH but not EYFP labeling, and white arrowheads show cell bodies with EYFP but not TH labeling. Abbreviations: fr – fasciculus retroflexus; IF – interfascicular nucleus; IPF – interpenducular fossa; IPR – interpeduncular nucleus; ml – medial lemniscus; mp – mammilary peduncle; MT – medial terminal nucleus; mtg – mammilotegmental tract; PBP – parabrachial pigmented nucleus, VTA; PN – paranigral nucleus, VTA; Rli – rostral linear nucleus; SNc – substantia nigra pars compacta; SNr – substantia nigra pars reticulata. **C**. Summary of targeting efficiency (left: percentage of TH-positive neurons exhibiting viral-driven EYFP fluorescence) and targeting fidelity (right: percentage of EYFP-positive neurons with TH labeling). Data were obtained from 2 male and 2 female mice; 6 sections containing the VTA were evaluated per mouse. In total, more than 10,000 TH-positive cells were evaluated for EYFP fluorescence and more than 10,000 EYFP-positive cells were evaluated for TH labeling. **D**. Representative traces showing action potentials induced by optical stimulation (1, 5, 10, 25 Hz) in VTA neurons expressing ChR2, but not VTA neurons expressing EYFP; scale: 20 mV/1 s. **E**. Summary of optically induced spike fidelity, or the percentage of optical stimuli that evoked a single action potential, as a function of stimulation frequency in ChR2-expressing VTA neurons (n=8 recordings from 2 male and 3 female mice).

To test whether the dual-virus approach permits optogenetic regulation of VTA DA neurons, C57BL/6J mice received an intra-VTA infusion of a 1:1 mixture of AAV8-mTH-Cre and an AAV harboring a Cre-dependent expression cassette for a channelrhodopsin/EFYP fusion protein (AAV8-EF1α-DIO-ChR2(H134R)-EYFP). After a 2-4-wk incubation period, we then employed slice electrophysiological approaches to characterize ChR2-expressing (EYFP-positive) VTA neurons in the PBP. ChR2-expressing neurons exhibited relatively large apparent capacitances (69±6 pF), low input resistances (267±31 MΩ), and large I_h_ currents (139±37 pA). Most (6/8; 75%) exhibited spontaneous activity (2.1±0.7 Hz) and long action potential durations (3.0±0.3 ms). These physiological properties are consistent with those we have reported for VTA DA neurons identified in prior studies using DAT-Cre mice [33-36]. Consistent with findings from studies involving TH-Cre or DAT-Cre lines [7,11], ChR2-expressing VTA neurons in C57BL/6J mice responded with high fidelity to brief pulses (5 ms) of blue light delivered at frequencies up to 25 Hz (**Fig. 1D,E**). In contrast, control (EYFP-expressing) VTA neurons did not respond to optical stimulation. Thus, the AAV-only approach permits optogenetic regulation of VTA DA neurons in C57BL/6J mice.

### Optical self-stimulation in C57BL/6J mice

We next tested whether AAV-mediated ChR2 expression in VTA DA neurons would support optical self-stimulation (OSS) in C57BL/6J mice (**Fig. 2A**). Control (EYFP-expressing) C57BL/6J mice, as well as DAT-Cre mice treated with the Cre-dependent ChR2 expression vector, were evaluated in parallel. Mice began acquisition sessions 16-19 d after viral infusion and fiber optic light guide implantation in the VTA; conditions were modeled after those in a study involving DAT-Cre mice [37] (**Fig. 2B**). Acquisition sessions (1 h/d) were conducted over 11 d using a fixed-ratio 1 (FR1) schedule of reinforcement. During these sessions, a nosepoke in the active port resulted in a 3-s optical stimulus delivered to the VTA at 50 Hz, with a 5-ms pulse duration.

**Figure 2.**
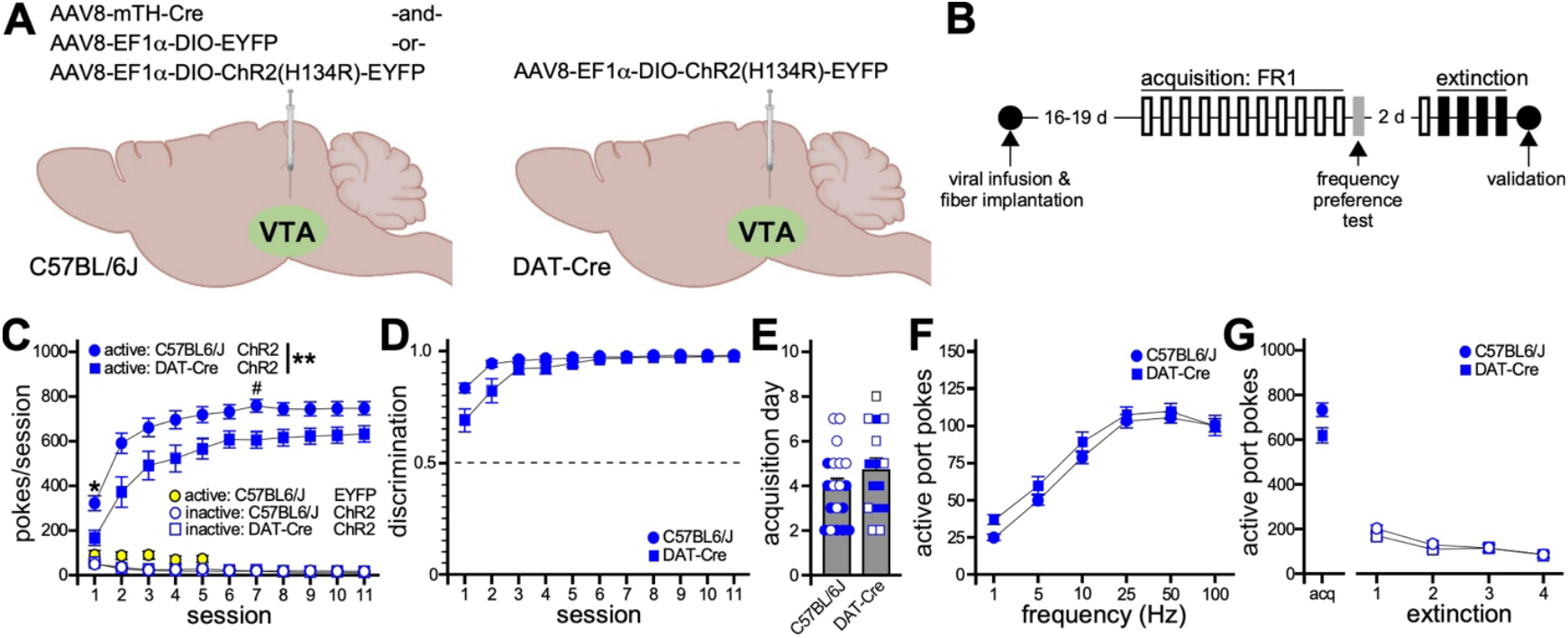
Optical self-stimulation in C57BL/6J mice. **A**. C57BL/6J mice were given intra-VTA infusions of AAV8-mTH-Cre and either AAV8-EF1α-DIO-ChR2(H134R)-EYFP or AAV8-EF1α-DIO-EYFP. DAT-Cre mice were given intra-VTA infusion of AAV8-EF1α-DIO-ChR2(H134R)-EYFP. **B**. Following viral infusion and fiber implantation surgery, subjects recovered for 16-19 d before beginning daily (1 h) optical self-stimulation acquisition sessions using an FR1 schedule of reinforcement; acquisition was conducted over an 11-d period. On day 12, subjects were evaluated in an optical frequency preference test. Following a 2-d rest period and 1 additional FR1 acquisition session, subjects began a 4-d extinction test. The accuracy of viral targeting and fiber placement was then evaluated in all subjects. **C**. Active (filled symbols) and inactive (open symbols) port nosepokes during acquisition sessions, for C57BL/6J (n=19) and DAT-Cre (n=15) mice expressing ChR2 in VTA DA neurons. Control C57BL/6J mice expressing EYFP in VTA DA neurons (yellow circles; n=6) exhibited low-level responding on the active port over 5 sessions. Two-way repeated measures ANOVA of active-port nosepoke data for ChR2-expressing C57BL/6J and DAT-Cre groups revealed a main effect of session (F_2.63,84.25_=108.5, p<0.0001) and genotype (F_1,32_=8.54, **p=0.0063), but no session x genotype interaction (F_2.63,84.25_=1.47, p=0.2327). Symbol: ^#^p<0.05 (within session). **D**. Discrimination index during acquisition sessions for C57BL/6J and DAT-Cre mice expressing ChR2 in VTA DA neurons. Two-way repeated measures ANOVA of discrimination index data revealed a main effect of session (F_1.59,50.95_=46.15, p<0.0001) but not genotype (F_1,32_=3.35, p=0.0766), and a session x genotype interaction (F_1.59,50.95_=6.63, p=0.0051). Symbols: *p<0.05 (within session). Pairwise comparison did not detect any significant differences between genotypes for any acquisition session. **E**. Acquisition day for C57BL/6J and DAT-Cre mice (t_1.25_=27.51, p=0.22; Student’s t-test with Welch’s correction). Individual data points for male and female subjects are denoted in open (female) and filled (male) symbols. **F**. Optical frequency test for C57BL/6J and DAT-Cre mice. For each animal, the number of active-port nosepokes measured during each 15-min frequency session was normalized to the number of active-port nosepokes measured at the frequency that evoked maximal responding. Two-way repeated measures ANOVA of frequency test data revealed a main effect of frequency (F_3.04,97.17_=249.0, p<0.0001) but not genotype (F_1,32_=1.45, p=0.24). Moreover, no frequency x genotype interaction was detected (F_5,160_=1.39, p=0.23). **G**. Extinction test for C57BL/6J and DAT-Cre mice. The number of active-port nosepokes recorded during the preceding acquisition session is shown for comparison only. Two-way repeated measures ANOVA of active-port nosepoke data during the 4 extinction sessions revealed a main effect of session (F_2.25,71.92_=36.4, p<0.0001) but not genotype (F_1,32_=1.00, p=0.32). Moreover, no frequency x genotype interaction was detected (F_2.25,71.92_=1.59; p=0.21).

While EYFP-expressing C57BL/6J control mice failed to develop significant active port responding over 5 sessions, all ChR2-expressing mice in the C57BL/6J and DAT-Cre groups developed robust and stable active-port responding (**Fig. 2C**), with a high degree of active:inactive port discrimination (**Fig. 2D**), over this same period. Interestingly, ChR2-expressing C57BL/6J mice exhibited higher levels of active-port responding than DAT-Cre mice. While ChR2-expressing C57BL/6J mice reached stable and high levels of active-port discrimination faster than DAT-Cre mice (**Fig. 2D**), the rate of acquisition – defined as the first of 3 consecutive sessions during which active port responding did not vary by more than 10% and discrimination index was >0.75 – did not differ significantly between genotypes (**Fig. 2E**).

Following the acquisition sessions, active-port responding as a function of optical stimulation frequency was evaluated in all subjects. Active-port responding was frequency-dependent for ChR2-expressing C57BL/6J and DAT-Cre mice. Maximal responding in both groups was seen between 25-100 Hz, and no genotype-dependent differences in sensitivity to optical stimulation frequency were detected (**Fig. 2F**). Finally, we evaluated rate of extinction of active-port responding over 4 sessions wherein a nosepoke in either port had no programmed response. ChR2-expressing C57BL/6J and DAT-Cre mice rapidly reduced active-port responding during the extinction sessions, with no evident genotype-dependent differences in rate or extent of extinction (**Fig. 2G**).

## DISCUSSION

Optogenetic manipulation of VTA DA neurons is a powerful strategy to investigate the relevance of VTA DA neurons and associated microcircuits to motivation, reinforcement learning, and addiction-like behavior. Studies involving optogenetic manipulation of VTA DA neurons have relied on transgenic DAT-Cre and TH-Cre rodent strains to confer neuronal specificity. As discussed above, there are caveats associated with available Cre driver lines [19,20]. TH-Cre knock-in mice, for example, may confer access to some neuron populations within and surrounding the VTA that lack detectable TH expression [19,21]. DAT-Cre mice appear to permit more selective targeting of VTA DA neurons [19], but may not afford access to critical VTA DA neuron sub-populations that make important contributions to reward-related behaviors [22]. In addition, reliance on transgenic Cre driver lines for studies of VTA DA neurons incurs costs associated with acquiring and in-house breeding of mutant strains.

The goal of this study was to develop an efficient approach to VTA DA neuron manipulation that can be used to support OSS studies and does not rely on transgenic Cre driver lines. Our strategy employs 2 AAV vectors to promote Cre-dependent transgene expression in VTA DA neurons, with selectivity for DA neurons achieved via a 2571 bp fragment of the mTH promoter [29]. This promoter was derived based on alignment with the rat TH upstream promoter that drives selective transgene expression in catecholaminergic neurons, including DA neurons of the substantia nigra and VTA [38,39]. The mTH promoter was shown to drive high efficiency (86%) and high fidelity (81%) reporter expression in TH-positive neurons in the mouse VTA [29].

Our assessment of efficiency (82%) and fidelity (73%) in targeting TH-positive VTA neurons in C57BL/6J mice using the dual-virus approach align well with published findings from studies employing mouse and rat TH promoters. However, targeting fidelity metrics in studies involving intra-VTA infusion of Cre-dependent AAV vectors in TH-Cre knock-in mice are substantially higher (98+%) [7,21]. This discrepancy may reflect limitations associated with use of promoter fragments to drive selective transgene expression, suggesting that elements outside of the core TH promoter are critical for ensuring neuronal selectivity. Viral load (titer and infusion volume) can also affect targeting fidelity when using AAV vectors with cell-specific promoters [40-42]. Efficiency and fidelity metrics related to VTA DA neuron targeting can also be impacted by the VTA sub-region(s) selected for quantitative analysis and regional variability in TH expression in the rodent VTA [43,44]. In addition, some VTA neurons express TH mRNA but do not produce TH protein [45], which would lower the assessment of targeting fidelity as defined by the fraction of neurons expressing viral transgene that also exhibit TH immunolabeling.

Using the AAV-only strategy to manipulate VTA DA neurons, we show that ChR2 in VTA DA neurons supports robust OSS behavior in C57BL/6J mice. Moreover, parallel comparison of ChR2-expressing C57BL/6J and DAT-Cre mice revealed greater active-port responding among subjects in the AAV-only cohort. Enhanced active-port responding may reflect the fact that TH-driven transgene expression provides access to more VTA DA neuron sub-populations than transgene expression driven by the DAT promoter [22-25]. Indeed, DAT expression is low or undetectable in TH-positive VTA neurons that project to the mPFC, NAc core and medial shell, and basolateral amygdala [22]. Moreover, several discrete VTA DA projections, including those targeting the NAc core or shell and the infralimbic region of the mPFC, can independently support optical self-stimulation in mice [13,14]. Lower responding levels in the hemizygous DAT-Cre mice used in this study may also be attributable to the reported lower DAT expression and function in the mesocorticolimbic system in this mutant strain [18,26]. Consistent with this premise, hemizygous DAT-Cre mice on a C57BL/6J background exhibited increased basal locomotor activity and decreased sensitivity to the locomotor-stimulatory effects of amphetamine, whereas these behaviors were normal in hemizygous TH-Cre mice on a C57BL/6J background [27].

The AAV-only approach described herein allows for manipulation of VTA DA neurons in support of OSS studies without reliance on existing transgenic Cre driver mouse lines. Potential applications for this approach include comparative assessments of reward circuit function across inbred mouse strains and exploration of genetic influences on reward circuit function using knockout and transgenic mouse lines.

## METHODS

### Animals

C57BL/6J mice (stock 000664) were purchased from The Jackson Laboratories (Bar Harbor, ME) and allowed to acclimate for at least 1 wk prior to experimental handling. DAT-Cre (B6.SJL*-Slc6a3*^*tm1*.*1(cre)Bkmn*^/J; stock 006660; RRID:IMSR_JAX:006660) mice were purchased from The Jackson Laboratories and were crossed with C57BL/6J mice to generate the DAT-Cre hemizygous offspring used in this study. Mice were group-housed and maintained on a 12 h light/dark cycle, and they had *ad libitum* access to food and water. All animal use was reviewed and approved by the Institutional Animal Care and Use Committee of the University of Minnesota. All experiments involving animals were designed, performed, and reported in accordance with institutional regulations and ARRIVE Guidelines (2.0) [46].

### Reagents

All viral vectors (>1x10^12^ genocopies/mL) were packaged using the AAV8 serotype and purified by the University of Minnesota Viral Innovation Core (VIC; Minneapolis, MN). pAAV-mTH-Cre plasmid was generated by the VIC using standard cloning techniques and pAAV-mTH-GFP (RRID:Addgene_99128; a gift from Viviana Gradinaru) as the backbone [29]. The pAAV-EF1α-DIO-hChR2(H134R)-EYFP plasmid was a gift from Karl Deisseroth (RRID:Addgene_20298) and was used as the backbone to generate pAAV-EF1α-DIO-EYFP.

### Intracranial manipulations

Procedures for intra-VTA infusion of virus in mice have been described previously [34,47]. Briefly, mice (6-10 wk) were anesthetized with isoflurane and placed in a stereotaxic frame (David Kopf Instruments; Tujunga, CA). A microinjector was lowered through a burr hole in the skull and used to deliver a unilateral infusion (0.4 μL /infusion at 0.1 μL/min) of virus (from bregma, AP: -2.75 mm, ML: ±0.55 mm, DV: -5.00 mm). For mice used in optical self-stimulation studies, a small screw was placed in the skull (from bregma: AP: -1.25 mm, ML: ±2.00 mm) prior to viral infusion to aid in stabilization of a dental cement cap used to close the surgical site and hold the fiber optic light guide in place. After viral infusion, a fiber optic light guide implant (400 um – NA0.66, 5mm; Doric Lenses Inc; Quebec, QC) was positioned 250 μm above the viral infusion site (from bregma, AP: -2.75 mm, ML: ±0.55 mm, DV: -4.75 mm). Unilateral viral infusion and fiber implantation were counterbalanced by hemisphere within each group. Mice recovered for 2-4 wk after surgery before processing for immunohistochemical or electrophysiological analysis, and 16-19 d before beginning behavioral testing.

### Immunohistochemistry

Brains were harvested and drop-fixed in 4% paraformaldehyde in phosphate buffered saline (PBS) at 4°C overnight, rinsed and equilibrated in a 30% sucrose solution at 4°C overnight, and then flash-frozen in an isopentane/dry ice bath. Six coronal sections (50 μm) containing the VTA were obtained per mouse by cryostat and collected in PBS in a 24-well plate. Free-floating sections were permeabilized in PBS containing 0.5% Triton X-100 and 2% donkey serum for 10 min and then transferred to blocking solution (PBS containing 0.1% Triton X-100 and 2% donkey serum) for 30 min at room temperature. Sections were incubated at 4°C overnight in blocking solution with 1:500 dilutions of primary polyclonal antibodies from Sigma-Aldrich (Burlington, MA) targeting TH (AB152, rabbit anti-TH; RRID: AB_390204) and from Abcam (Waltham, MA) targeting GFP (AB6673, goat anti-GFP; RRID: AB_305643). Sections were then incubated in 1:500 dilutions of Cy™3 donkey anti-rabbit (711-165-152; RRID: AB_2307443) and Alexa Fluor® 488 donkey anti-goat (705-405-003; RRID: AB_2340428) secondary antibodies from Jackson ImmunoResearch Laboratories, Inc. (West Grove, PA) for 2 h at room temperature. Sections were then mounted using ProLong™ Gold Antifade Mountant with DNA stain DAPI (P36941, Invitrogen; Chicago, IL) and imaged using a BZ-X810 fluorescence microscope (Keyence; Itasca, IL).

TH and EYFP expression (and their overlap) were quantified in 6 sections containing the VTA per mouse, from 2 male and 2 female mice, using CellProfiler™ (RRID:SCR_007358) [48] and a modified version of a published analysis pipeline [49]. In brief, CellProfiler™ was used to subtract background fluorescence and segment cells based on nuclear (DAPI) staining in each section, within the region of viral infusion (defined by the scope of EYFP expression). Initial cell segmentation was expanded to include somatic fluorescent labeling to identify expression-positive cells. The probability that a cell was positive for TH or EYFP was calculated using a binomial distribution test comparing the area of expression within a cell against the area of background fluorescence. Data from the 6 sections were then aggregated and organized using a Python script (GitHub 10.5281/zenodo.18511410) to calculate targeting efficiency and fidelity for each mouse, based on the number of cells positive for 1 or both targets. Final targeting efficiency and fidelity metrics are derived from analysis of more than 10,000 TH-positive and 10,000 EYFP-positive cells.

### Slice electrophysiology

Horizontal slices (225 μm) containing the VTA were prepared as described [33]. Whole-cell recordings from EYFP-positive cells in the VTA were acquired in an oxygenated aCSF bath at 33-35ºC, using a Multiclamp 700B amplifier and pCLAMPv.10.6 software (Molecular Devices, LLC; San Diego, CA), as described [35]. After whole-cell access was achieved, I_h_ current was measured in voltage-clamp mode (V_hold_ = -45 mV) using a 200-ms voltage step to −120 mV, and spontaneous activity was monitored for 1-min in current-clamp mode (I = 0). Optically evoked action potentials were measured in current-clamp mode (I=0) using 5 ms pulses of 470 nm wavelength light (2-5 mW), delivered for 3 s at 1, 5, 10, 25 Hz through a 40x objective. All measured and command potentials factored in a liquid junction potential of −15 mV.

### Optogenetic self-stimulation (OSS)

Operant conditioning boxes constructed with ARDUINO boards (Arduino Uno or Elegoo Uno R3) were built and controlled as described [37]. Each box has 2 nosepoke ports and green LED cue lights mounted above the ports. Prior to each OSS session, mice were allowed at least 30 min to acclimate to the testing room. Mice were then hand-restrained and a fiber optic patchcord (MFP 400/430/1100-NA0.57 – 1m) from Doric Lenses Inc. (Quebec, Canada) was connected to the fiber optic light guide implant. The fiber optic patchcord was connected to a fiber-coupled LED (M470F4) from Thorlabs, Inc. (Newton, NJ) through a fiber optic rotary joint (FRJ 1x1 400/430/LWMJ-0.57; Doric Lenses Inc.). During each 60-min acquisition session, a nosepoke in the active port resulted in illumination of the green cue light above the port and concurrent 3-s optical stimulus (LED 470 nm/20 mW). The optical stimulus for acquisition sessions was delivered at 50 Hz, with a 5-ms pulse duration. A nosepoke in the inactive port resulted only in illumination of the green cue light above the port. Discrimination index was defined as the number of active-port nosepokes divided by the total number of nosepokes during a session. Acquisition of OSS was defined as the first of 3 consecutive days when the number of active-port nosepokes varied by less than 10% and discrimination index exceeded 0.75.

The stimulation frequency preference test was conducted 1 d after mice finished the 11 acquisition sessions. During this 90-min test, the frequency of optical stimulus elicited by an active-port nosepoke was changed every 15 min (900 s) in the following order: 100 Hz, 50 Hz, 25 Hz, 10 Hz, 5 Hz, 1 Hz. The number of active-port nosepokes was measured at each frequency for each subject. These values were then normalized to the number of active-port nosepokes measured at the stimulation frequency that generated the highest level of responding by that mouse. Extinction of OSS behavior was assessed over 4 consecutive days in 60-min sessions during which responding in active and inactive ports had no programmed consequences.

Following extinction, brains were harvested and 225 μm horizontal sections containing the VTA were obtained by vibratome. Brightfield and fluorescent images were acquired using a BZ-X810 fluorescence microscope (Keyence). Light guide placement and the scope of AAV-driven EYFP fluorescence were evaluated using the Allen Mouse Brain Atlas as a reference [50]. Data from mice with mistargeted light guide implants, or from mice that lacked fluorescence or exhibited significant fluorescence outside of the VTA (*e*.*g*., in the adjacent substantia nigra pars compacta), were excluded from analysis.

## Data analysis

Data were analyzed using Prism v.10.0.2 software (GraphPad Software; Boston, MA). All studies included balanced numbers of male and female mice, and data were evaluated first for sex differences. We did not observe any effect of sex or interaction involving sex for any measured endpoint in this study. As such, data from male and female mice were pooled and are presented throughout as mean ± SEM. Pooled behavioral data were evaluated for session and/or genotype effects using Student’s t test or 2-way repeated measures ANOVA. Sidák’s multiple comparisons test was used for pairwise comparisons if warranted. Differences were considered significant if p<0.05.

## Acknowledgements

The authors thank members of the Wickman lab (McKinzie Frederick, Anna Souders) for care of the mouse colony and technical support, and Dr. Sarah Mulloy for providing guidance related to the analysis pipeline for the immunohistochemistry approach. This study was supported by the following NIH awards: R01 AA027544 and R01 DA034696 (KW), R25 DA057802 (JCB), F31 DA062412 (ALW), T32 DA007234 (ALW), and T32 NS105604 (EHM). Financial support for viral vector production came from the Viral Innovation Core, part of the University of Minnesota *Center for Neural Circuits in Addiction* (P30 DA048742).

## Author contributions

Ezequiel Marron Fernandez de Velasco: conceptualization, data curation, formal analysis, investigation, methodology, resources, supervision, validation, visualization, writing – review & editing

John C. Brent IV: data curation, formal analysis, investigation, methodology, visualization, writing – review & editing

Alex L. Welter: formal analysis, investigation, methodology, validation, visualization, writing – review & editing

Praseuth Yang: formal analysis, investigation, methodology, validation, visualization, writing – review & editing

Annelise Wickman: methodology, resources, writing – review & editing

Eric H. Mitten: methodology, writing – review & editing

Kevin Wickman: conceptualization, funding acquisition, project administration, supervision, validation, visualization, writing-original draft, writing – review & editing

## Data availability

The datasets generated during and/or analyzed during the current study are available from the corresponding author on reasonable request.

